# Viral genome wide association study identifies novel hepatitis C virus polymorphisms associated with sofosbuvir treatment failure

**DOI:** 10.1101/2020.05.05.077230

**Authors:** David A Smith, Andrea Magri, Rory Bowden, Nimisha Chaturvedi, Jacques Fellay, John McLauchlan, Graham R. Foster, William L Irving, STOP-HCV Consortium, Peter Simmonds, Vincent Pedergnana, Eleanor Barnes, M. Azim Ansari

**Affiliations:** Peter Medawar Building for Pathogens Research, Nuffield Department of Medicine, University of Oxford, Oxford OX1 4BH, UK; Wellcome Trust Centre for Human Genetics, University of Oxford, Oxford, OX1 4BH, UK; School of Life Sciences, École Polytechnique Fédérale de Lausanne, Lausanne, Switzerland; Precision Medicine Unit, University Hospital and University of Lausanne, Lausanne, Switzerland; Swiss Institute of Bioinformatics, Lausanne, Switzerland; MRC-University of Glasgow Centre for Virus Research, Glasgow, UK; Barts Liver Centre, Blizard Institute, Queen Mary University of London, London, UK; University of Nottingham and Nottingham University Hospitals NHS Trust; French National Centre for Scientific Research (CNRS), Laboratory MIVEGEC (CNRS, IRD, UM), Montpellier, France

**Keywords:** HCV, Hepatitis C Virus, Genotype 3, Direct Acting Antivirals, DAA, Sofosbuvir

## Abstract

Chronic hepatitis C virus (HCV) infection is a major cause of chronic liver disease, cirrhosis and hepatocellular carcinoma worldwide. With the recent development of direct acting antivirals (DAA), treatment of chronically infected patients has become highly effective although a subset of patients do not respond to therapy. Sofosbuvir is a common component of current de novo or salvage combination therapies. We used pre-treatment whole genome sequencing of HCV from 507 patients infected with HCV subtype 3a and treated with sofosbuvir containing regimens to detect viral polymorphisms associated with response to treatment. We found that three common polymorphisms present in HCV NS2 and NS3 proteins (not direct targets of sofosbuvir) were associated with reduced treatment response. These polymorphisms were enriched in post-treatment HCV sequences of patients unresponsive to treatment; they were also associated with lower reductions in viral load in the first week of therapy. The finding of polymorphisms in NS2 and NS3 proteins associated with poor treatment outcomes emphasises the value of more systematic genome-wide analyses of HCV in uncovering indirect but clinically relevant mechanisms of antiviral resistance.

## Introduction

An estimated 70 million people worldwide are chronically infected with hepatitis C virus (HCV), a major aetiology of chronic liver disease which can culminate in cirrhosis and hepatocellular carcinoma^1^. Since 2011, the rapid development of oral direct-acting antivirals (DAA) which target different HCV proteins has resulted in significant improvements in the safety and efficacy of treatments that cure HCV infection (sustained virologic response (SVR)). Combinations of DAAs targeting different HCV proteins regularly achieve SVR rates in excess of 95%^2,3^.

Sofosbuvir is a nucleotide analogue and competitively binds to and blocks the HCV polymerase (coded by non-structural 5B (NS5B) gene) and inhibits viral replication^4,5^. Sofosbuvir is a key component in several currently utilised treatment regimens and is recommended for use in combination with other DAAs such as NS5A inhibitors ledipasvir, daclatasvir or velpatasvir and the NS3 protease inhibitor voxilaprevir. Sofosbuvir in combination with velpatasvir and voxilaprevir has become one of the recommended salvage regimens for patients unresponsive to previous DAA therapies^6,7^. Sofosbuvir is effective against all HCV genotypes and has a high barrier to the development of resistance^8^. However, pre-treatment resistance associated substitutions (RAS) for both NS3 and NS5A inhibitors used in combination with sofosbuvir^9–12^ are commonly reported. Subtypes 3b and 4r have been shown to be inherently resistant to current NS5A inhibitors^13^. Where RAS are present, they can cause a large reduction in the potency of NS3 and NS5A inhibitors, making sofosbuvir the main active compound in these regimens.

Only a small number of sofosbuvir-associated RAS have been reported^8,14^ relative to other DAAs. In multiple clinical trials using sofosbuvir, NS5B substitutions L159F, S282T, C316H/N, L320F and V321A are reported as RAS and have been detected post-treatment in patients who do not achieve SVR^14^ (in a small subset of those patients). The acquisition of these RAS has been shown to have a large fitness cost for the virus *in vitro*^11^ and thus they rarely persist for long after treatment failure^15^. An exception is the S282T RAS, which persists if compensatory mutations are also acquired^16^. Certain HCV subtypes (4r) have been shown to acquire this substitution more readily^9,10^. Furthermore, with the exception of S282T, all sofosbuvir RAS show negligible increases in resistance using *in vitro* models^11^ and patients who were unresponsive to sofosbuvir containing regimens, often do not carry any of these RAS at baseline or post-treatment. We recently reported the NS5B A150V polymorphism as conferring resistance to sofosbuvir^17^ using patient data and in vitro assays. This polymorphism unlike previously reported substitutions is common in pre-treatment HCV subtype 3a sequences. To date, the mechanism of treatment failure in the small proportion of individuals who do not respond to sofosbuvir-based regimens remains unknown. Most studies of sofosbuvir RAS have focused on NS5B protein and the comparison of the post-treatment to baseline sequences which can only detect treatment emergent RAS. Investigating the impact of the pre-existing polymorphisms in NS5B and other HCV proteins could provide novel insight into viral determinants of sofosbuvir treatment and help us stratify patients more effectively.

In this study we used full length HCV genomes and deep sequencing data from 507 patients infected with HCV gt3a and treated with sofosbuvir DAA mono-therapy from the BOSON clinical trial^18^, to identify viral determinants of sofosbuvir treatment outcome. We performed a viral genome wide association study (V-GWAS) of baseline sequences and identified three viral polymorphisms outside the NS5B protein, which were significantly associated with SVR which we refer to as treatment outcome polymorphisms (TOPs). Additionally, we found a stepwise reduction in the SVR rate associated with an increase in total number of TOPs. The TOPs were also associated with a lower reduction in viral load during the first week of therapy and were enriched in the post-treatment sequences in patients who did not achieve SVR. The approach demonstrates the potential of genome-wide assessment of viral sequences to provide unexpected insight into molecular interactions and their importance in improving treatment of viral infections.

## Results

### Non-viral factors associated with SVR

Baseline whole genome HCV sequences were generated for 568 patients enrolled in the BOSON study^18,19^, 518 of which were gt3, 49 were gt2 and one was a gt1a/2b recombinant virus. HCV is a highly diverse pathogen and to limit the impact of virus population structure on our analysis, we restricted the analyses to samples from patients infected with subtype 3a (N=507). Overall 82% (416/507) of the gt3a patients achieved SVR.

To account for non-viral factors associated with SVR, we used logistic regression to test for association between treatment outcome and patient’s cirrhosis status, gender, baseline viral load, prior IFN-based treatment and *IFNL4* SNP rs12979860 genotypes (CC vs non-CC) in a multivariate model. Cirrhosis status was significantly associated with SVR (P=1.5×10^−4^), having the largest impact on treatment response (failure rate in cirrhotic patients 29%=47/163 and in non-cirrhotic patients 13%=44/344). Male gender and the non-CC genotype of the *IFNL4* SNP rs12979860 were also significantly associated with increase in treatment failure rate (P_gender_=0.016, P_*IFNL4*_=6.4×10^−3^). Previous IFN-based treatment and higher viral load were associated with statistically non-significant increases in treatment failure (**Supplementary Figure *1***). In all our subsequent regression-based analysis *INFL4* genotype, cirrhosis, gender, previous IFN-based treatment and log10 of baseline viral load were added as covariates to account for possible confounders.

**Figure 1:**
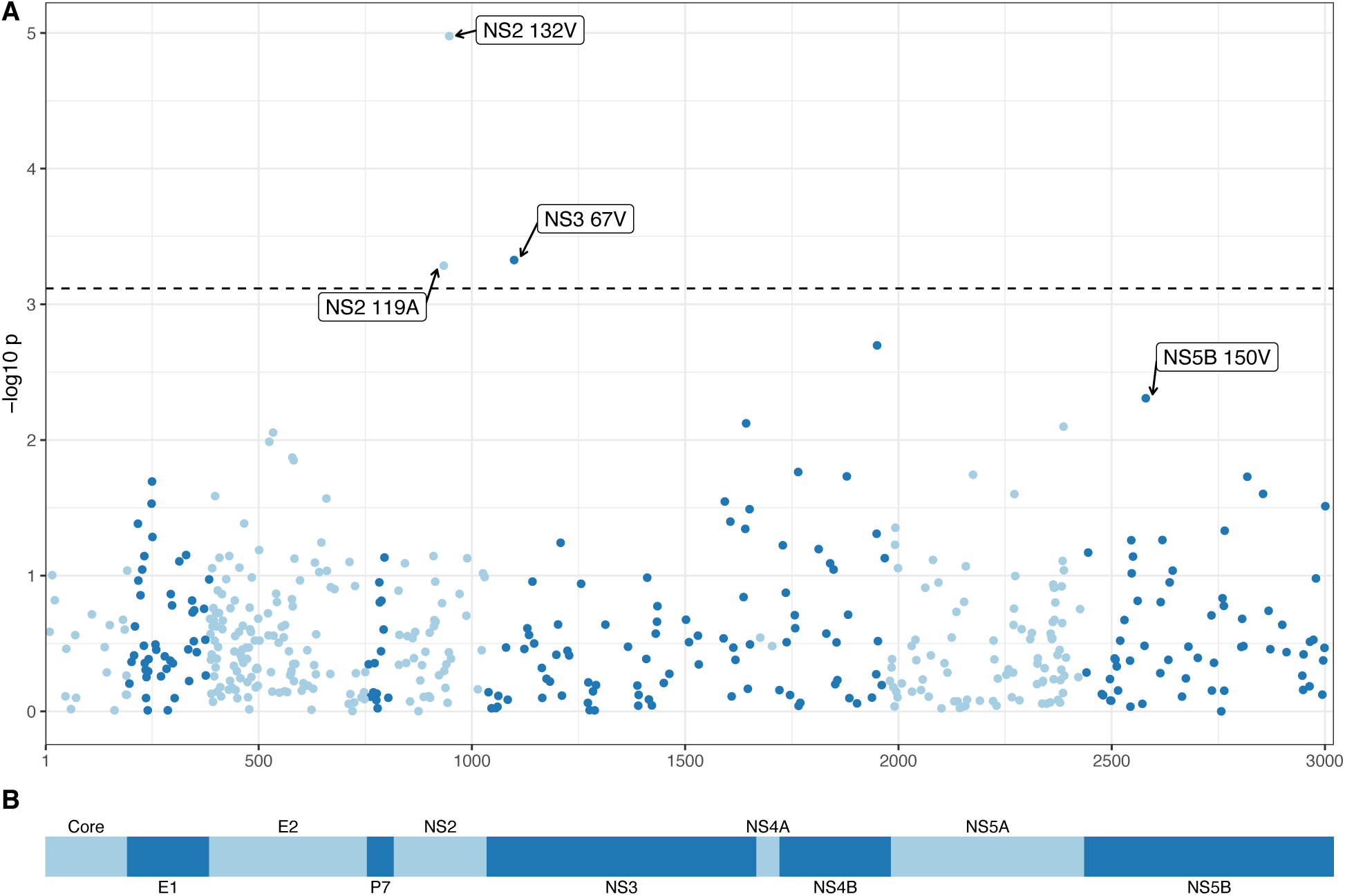
Association between HCV amino acids and sofosbuvir based treatment outcome. A) Manhattan plot of association tests between HCV amino acids and treatment outcome. At each viral site only the p-value for the most associated amino acid is plotted. The dashed line indicates 15% FDR. For the three sites significantly associated with outcome and the most associated site in the NS5B protein, the amino acid associated with the lowest cure rate at the site and its position within its respective protein is indicated. B) Schematic of HCV polyprotein.

During the BOSON clinical trial, patients were randomised into three treatment arms of sofosbuvir and ribavirin for 16 weeks (16 SOF+RBV), sofosbuvir and ribavirin for 24 weeks (24 SOF+RBV), or sofosbuvir plus peginterferon-alfa and ribavirin for 12 weeks (12 SOF+RBV+IFN). These treatment arms had different SVR rates, but as patients were randomised into these treatment arms (which means viruses were also randomised), the confounding between viral factors and the treatment arms is minimised. We therefore decided to not include treatment arm as a covariate in our analysis as it reduces the power of the tests but have investigated the association between TOPs and the treatment arms to ensure there is no correlation between the two.

### Previously reported sofosbuvir RAS

We investigated the baseline sequences for presence of previously reported sofosbuvir RAS in the NS5B protein (A150V, L159F, S282T, C316H/N, L320F and V321A)^8,14,20^. Only A150V (n=204) and L159F (n=1) were detected in the consensus sequences (sequence data for these sites was not available for six patients). A150V was significantly associated with outcome (P=4.92×10^−3^) and patients carrying valine at this site had an SVR rate of 75% (n=153/204) vs. 86% (n=257/297) for patients with other amino acids at this site. We have previously reported this association in a subset of this cohort and showed it reduces sensitivity to sofosbuvir using *in vitro* assays^17^. We have also previously reported an association between this viral site and host *IFNL4* SNP rs12979860^19^ genotypes. Stratifying the patients based on the *IFNL4* SNP rs12979860 genotypes (CC vs. non-CC,(genotyping data not available for two patients)), we observed that, among CC patients, the impact of valine on SVR rate was minimal (SVR_valine_ 85%=35/41, SVR_non-valine_ 87%=129/148, Fisher’s exact test P=0.80). On the contrary, among non-CC patients it reduced SVR rate substantially (SVR_valine_ 72% = 117/162, SVR_non-valine_ 85%=127/148, Fisher’s exact test P=0.0036). However, the interaction term was statistically nonsignificant (P=0.36).

Analysis of minor populations in the NGS data revealed very low frequencies (<1% of reads) of S282T (two patients), C316N (one patient), L320F (two patients) and V321A (eight patients).

### Identification of TOPs using viral genome wide association study (V-GWAS)

A major advantage of HCV whole genome sequencing is the possibility of testing viral genetic variants for associations with SVR across the entire viral genome to identify potentially novel associations between HCV genetic variants and response to sofosbuvir treatment. Initially, we investigated whether any of the HCV lineages were associated with treatment outcome. To do this we built a maximum likelihood phylogenetic tree (**Supplementary Figure 2**) from baseline consensus sequences. Analysis by treeBreaker software^21^ did not reveal any clades that had a different SVR rate from the rest of the tree (Bayes factor=0.969, see **Methods**).

We then performed a V-GWAS, using logistic regression to test for association between sofosbuvir treatment outcome and encoded amino acids at each site, including the non-viral factors previously indicated as covariates. The first three viral genetic principal components (PCs) and the first three host genetic PCs were also included as covariates to account for host-virus population co-structuring. In our analysis, SVR status was used as the response variable and the presence and absence of each amino acid as the explanatory variable. We only tested amino acids that were present in sequences from at least 20 individuals; this resulted in 1,010 tests at 484 sites. At a false discovery rate (FDR) of 15%, three viral polymorphisms were significantly associated with SVR (**Figure *1*** and **Supplementary Table 1**, the numbers below may not add to 507 as not all sites on the viral polyprotein were available for all samples, see **Supplementary Table 1**). The most significant association with SVR was at position 132 in NS2 protein (P=1.05×10^−5^), where isoleucine (80%=405/506) and valine (20%=100/506) alternated (one isolate carried leucine). Valine was associated with a reduction in treatment response (SVR 66%=66/100) relative to the isoleucine residue (86%=348/405). Two additional significant associations with reduction in SVR rate were valine at position 67 in the NS3 protein (P=4.71×10^−4^) and alanine at position 119 in the NS2 protein (P=5.18×10^−4^) (**Figure 1** and **Supplementary Table 2**). In the NS5B protein (the direct antiviral target of sofosbuvir), valine at position 150 was ranked first (lowest p-value, P=4.92×10^−3^) although the association was not significant genome-wide at a 15% FDR. For the remainder of this work NS5B A150V is included as a TOP.

Next we used an alternative chronic HCV infection cohort of 144 patients provided by HCV Research UK to replicate the findings. All patients had cirrhosis and 107 (75%) were treated with sofosbuvir in combination with an additional NS5A inhibitor such as daclatasvir, ledipasvir or velpatasvir, while the remaining 37 (25%) patients were treated either with SOF+RBV or SOF+RBV+INF. In this replication cohort, only NS2 119A was nominally associated with SVR (P=0.045). Overall, the effect size estimates were similar for three of the four TOPs (NS2 119A, NS2 132V and NS5B 150V), but they had larger confidence intervals due to smaller samples sizes and additional DAAs used in the therapy (**Supplementary Figure *3*A**).

### Covariation of TOPs and their impact on sustained virologic response

To understand the impact of presence of multiple TOPs in each sequence on SVR, we stratified patients based on the number of TOPs present in their baseline consensus sequences (**Figure *2*A**). The SVR rate for patients whose virus carried no TOPs was 95% (133/140) which reduced to 50% (13/26) for patients with three TOPs. Only one patient carried all four TOPs and this patient achieved SVR. The step-wise reduction in the SVR rate associated with the increase in the number of TOPs present at baseline was highly significant (logistic regression P=6.6×10^−10^).

**Figure 2:**
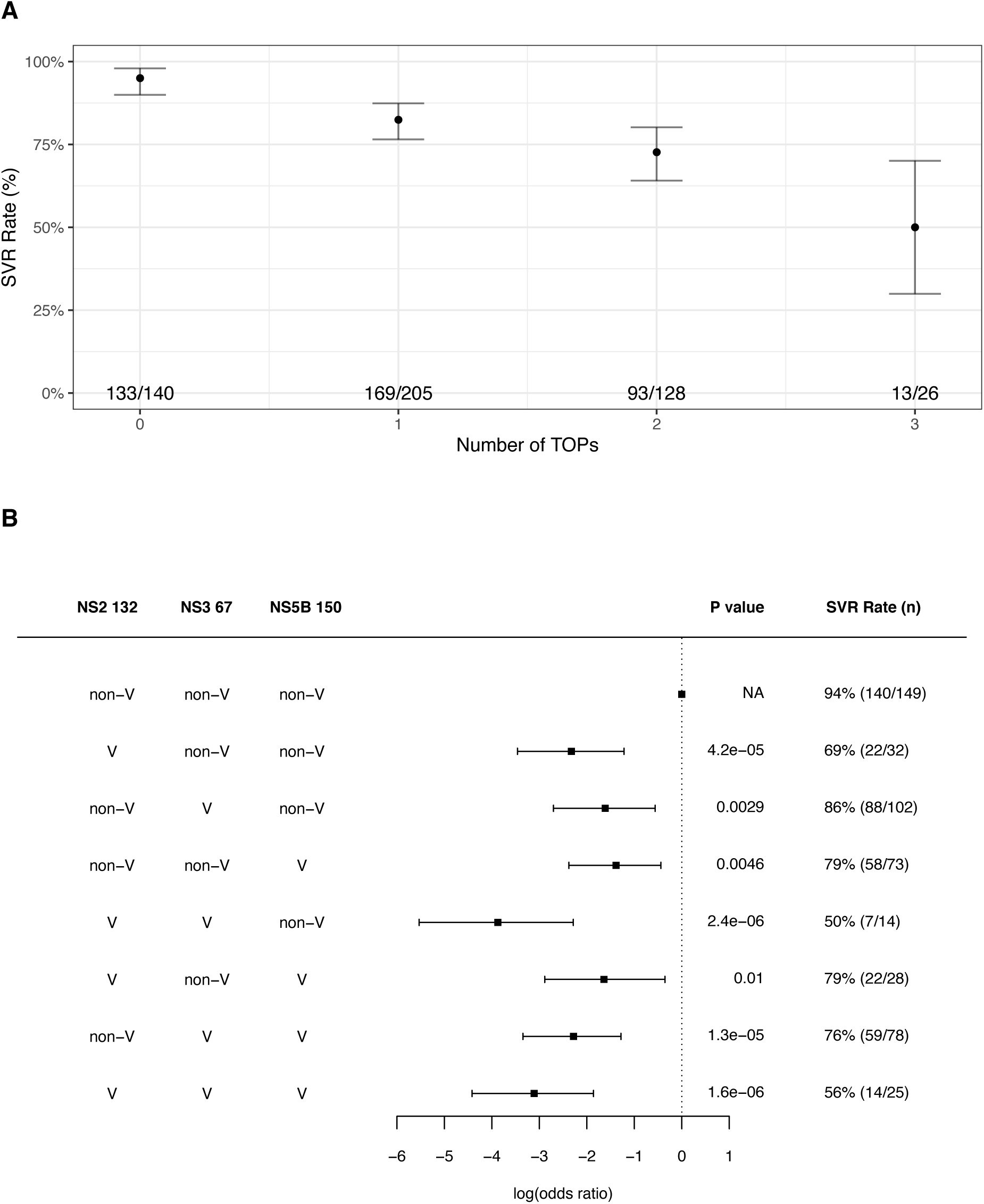
Covariation of TOPs and their impact on SVR rate. A) SVR rate for patients with different numbers of TOPs in baseline sequences. The dots indicate the SVR rate in each group and the lines indicate its 95% confidence intervals. The numbers in each group are shown at the bottom of the figure. B) Covariation between the 3 novel TOPs and their association with treatment outcome. Combinations tested are listed in the table on the left. The squares show the estimated effect size (log(OR)) for each group and the lines show its 95% confidence interval. The P value and the SVR rate for each group are shown on the right.

We also investigated the covariation between these TOPs and their impact on SVR. NS2 119A was excluded from this analysis because of its low frequency (4%=22/505). We tested for association (covariation) between the three pairs using logistic regression, including the first three viral PCs and host *IFNL4* SNP rs12979860 genotypes (CC vs. non-CC) as covariates to account for virus population structure and the impact of *IFNL4* on the HCV amino acids. NS5B 150V was significantly associated with NS2 132V (p =0.004) and nominally with NS3 67V (P=0.02). NS2 132V and NS3 67V were not significantly associated with each other (P=0.83). Stratifying the patients by presence and absence of TOPs (valine residue) at the three sites of NS2 132, NS3 67 and NS5B 150, we observed that patients carrying viruses with non-valine residues at all three sites had the highest SVR rate (94%=140/149), while carrying any one of these TOPs individually or in combination reduced SVR rate significantly (**Figure 2B**). Carrying valine at both NS2 132 and NS3 67 sites, but not NS5B 150 was associated with the lowest SVR rate of 50% (P=2.4×10^−6^). We also tested for non-additive interaction effects on SVR between the three sites and found that the only significant non-additive interaction effect was between NS2 132 and NS5B 150 sites (interaction P=0.01) where carrying valine at both sites resulted in SVR recovery (79%=22/28) rather than an additive reduction.

### TOPs effects on viral load reduction during first week of therapy

If these TOPs reduce the impact of treatment, one would expect to observe differences in the reduction in viral load during therapy between patients with and without these TOPs. Thus we evaluated the impact of these novel TOPs on the viral load reduction from baseline to week one of therapy and tested for difference in mean reduction between patients with and without each TOP and also their combination (**Figure *3***). At each of the sites, the mean reduction in viral load was smaller for TOPs relative to non-TOPs residues. The reduction in viral load was nominally significant for patients whose virus carried TOPs at sites NS2 119 (P=0.013), NS2 132 (P=0.018) or NS3 67 (P=0.029) relative to those whose virus did not carry the TOPs at these sites. The impact of the total number of TOPs carried by each patient on the viral load reduction was also significant (linear regression, P=0.006, **Figure *3*B**). Investigating the eight possible combinations of the three sites of NS2 132, NS3 67 and NS5B 150 we observed that the presence of valine at the two sites of NS2 132 and NS3 67 and non-valine at NS5B150 resulted in the lowest reduction in viral load during therapy (P=0.00091, **Supplementary Figure *4***).

**Figure 3:**
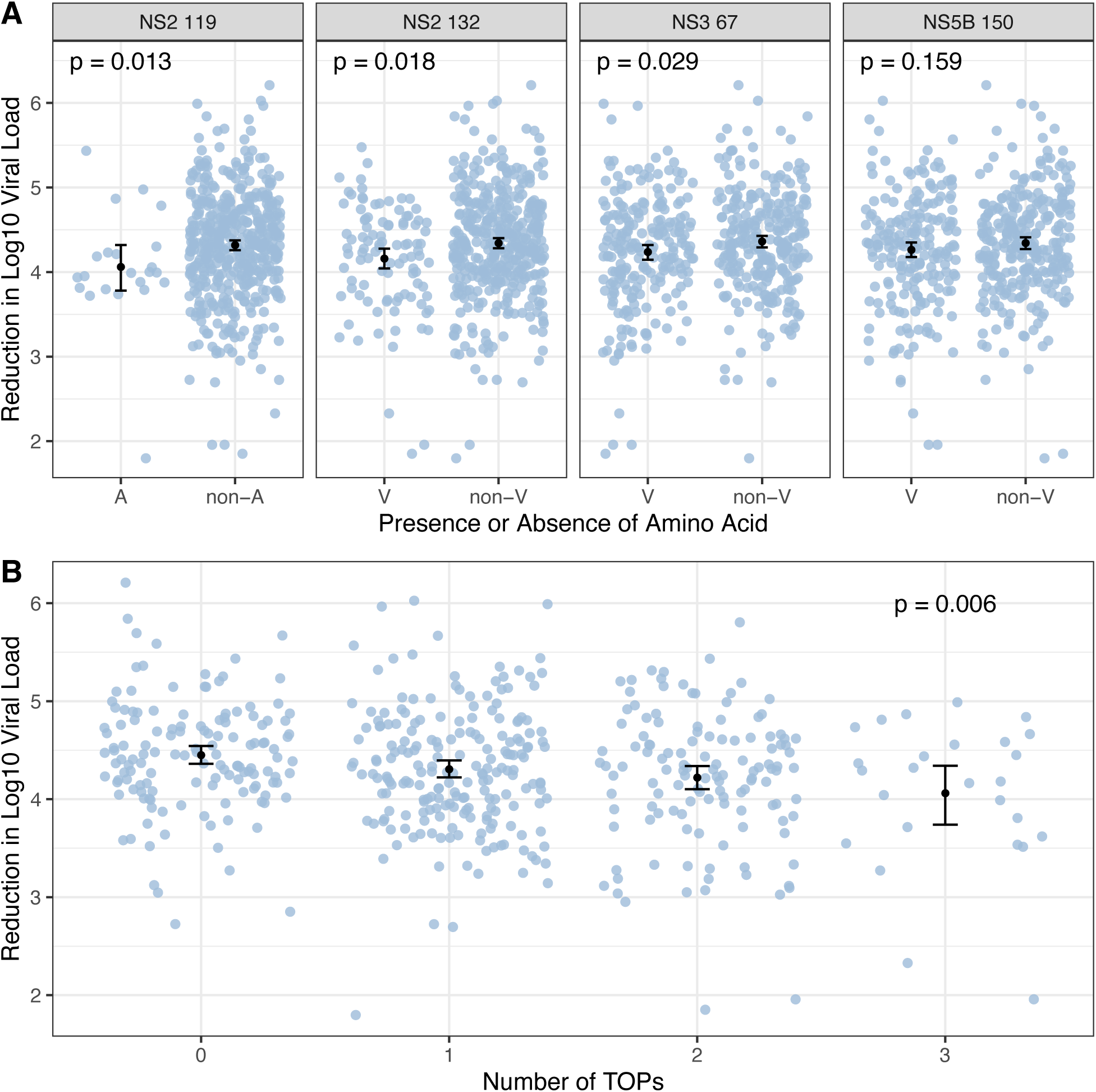
Reduction in viral load during the first week of therapy and its association with TOPs. A) Reduction in log10 viral load between baseline and week one of therapy for viruses with or without each TOP. The mean change in viral load is shown as a black dot for each TOP and the lines indicate its 95% confidence intervals. P values for difference in mean calculated using one-sided Mann-Whitney test. B) Reduction in log10 viral load between baseline and week one of therapy against increasing numbers of TOPs. The mean change in viral load is shown as a black dot and the lines indicate its 95% confidence interval. P value for association between number of TOPS and reduction in log10 viral load calculated using linear regression.

### Prevalence and enrichment of TOPs in post treatment sequences

Ninety-one patients in this trial did not achieve SVR and we were able to generate 73 whole genome consensus sequences from plasma samples taken at 12 weeks post-treatment (12WPT). These consensus sequences were scanned for the presence of previously reported sofosbuvir RASs. NS5B 159F was detected in four isolates. This represents a significant change in the distribution of this RAS in the post treatment sequences relative to the baseline sequences (binomial test, P=1.65×10^−5^). The NS5B RAS 282T, 316H/N, 320F and 321A were not detected in any of the post treatment consensus sequences. On analysis of the quasi-species data the 159F was detected in two additional 12WPT samples as minor variant (<15% of reads). The 321A was detected at low frequencies (<1% of reads) in two samples.

We also investigated the distribution of the TOPs identified using baseline sequences in our study in the 12WPT sequences. A binomial test was performed to identify changes in the distribution of the TOPs at these sites between the baseline population and the 12WPT sequences **(Table 1)**. NS5B 150V was significantly enriched in the 12WPT sequences (P=0.00088) and NS2 132V (P=0.019) and NS3 67V (P=0.023) were nominally significantly enriched. The majority of these polymorphisms were present at baseline and were carried forward to the post-treatment isolates suggesting that the increase in prevalence is due to enrichment of pre-existing polymorphisms rather than substitutions during therapy. However, this was not the case for the previously reported NS5B 159F RAS, where three of the four RAS had developed during therapy.

**Table 1:**
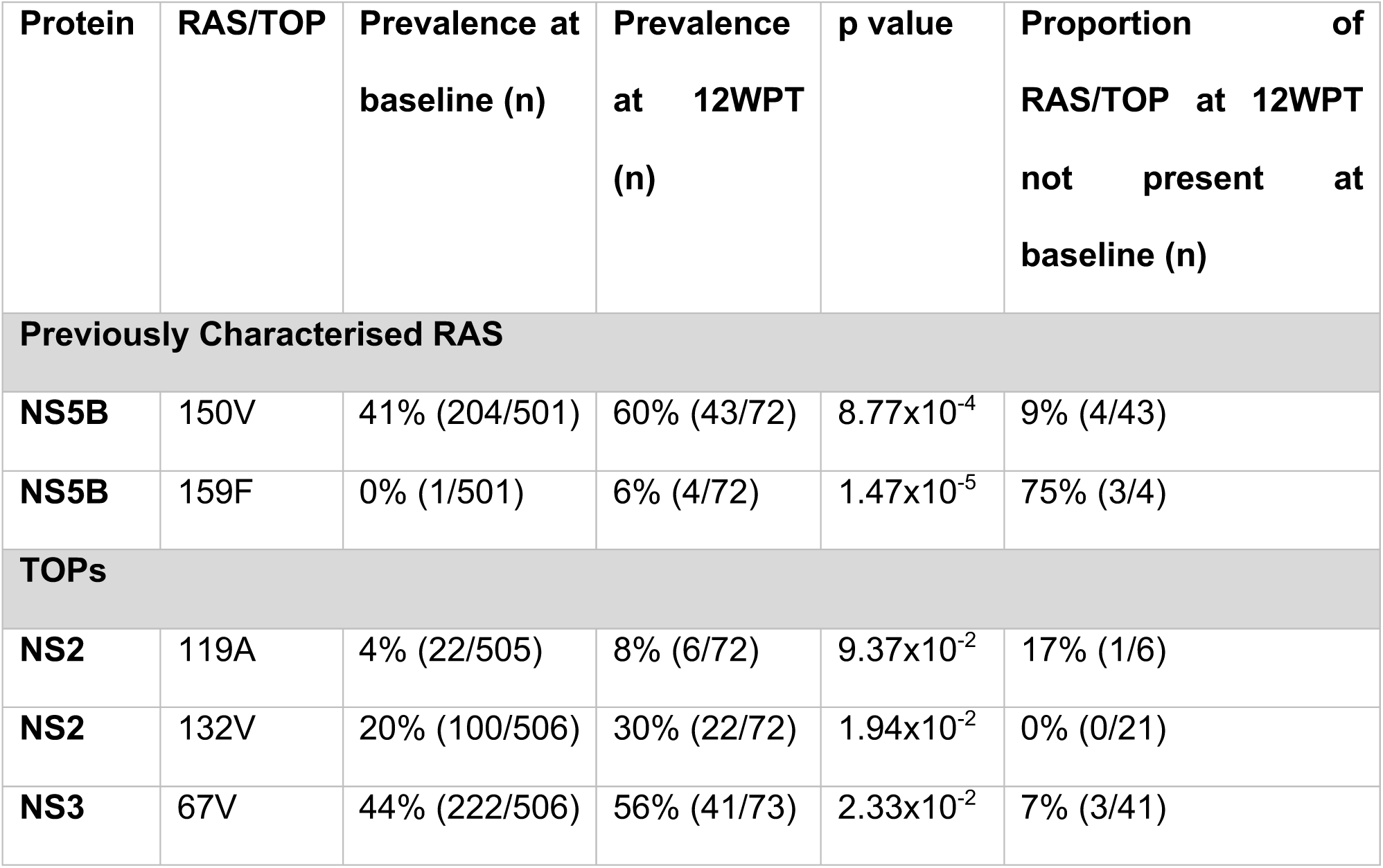
Prevalence of RAS and TOPs at baseline and 12 weeks post treatment. The percentage of the population with each RAS or TOP is shown at baseline and 12 weeks post treatment (12WPT). These distributions were compared using a one-sided binomial test and the p value is shown. The percentage of patients where the amino acid at baseline has changed to the RAS or TOP in post-treatment sequence is also shown.

| Protein | RAS/TOP | Prevalence at baseline (n) | Prevalence at 12WPT (n) | p value | Proportion of RAS/TOP at 12WPT not present at baseline (n) |
| --- | --- | --- | --- | --- | --- |
| **Previously Characterised RAS** |  |  |  |  |  |
| **NS5B** | 150V | 41% (204/501) | 60% (43/72) | $8.77 \times 10^{-4}$ | 9% (4/43) |
| **NS5B** | 159F | 0% (1/501) | 6% (4/72) | $1.47 \times 10^{-5}$ | 75% (3/4) |
| **TOPs** |  |  |  |  |  |
| **NS2** | 119A | 4% (22/505) | 8% (6/72) | $9.37 \times 10^{-2}$ | 17% (1/6) |
| **NS2** | 132V | 20% (100/506) | 30% (22/72) | $1.94 \times 10^{-2}$ | 0% (0/21) |
| **NS3** | 67V | 44% (222/506) | 56% (41/73) | $2.33 \times 10^{-2}$ | 7% (3/41) |

## Discussion

In this study we report the first systematic and comprehensive investigation of the viral determinants of treatment outcome in a large cohort of HCV gt3a infected patients treated with SOF+RBV+/-INF. We generated HCV whole genomes using next generation sequencing technologies for 507 patients at baseline and 73 patients who failed therapy at 12WPT. Using V-GWAS we performed a systematic assessment of the impact of HCV amino acids across all proteins on sofosbuvir treatment outcome. In this study we report for the first time, amino acid polymorphisms in HCV NS2 and NS3 proteins associated with treatment outcome. We also examined combinations of TOPs, the total number of TOPs carried, reduction in viral load during therapy and post-treatment failure sequences for additional evidence of the impact of these novel TOPs. We expect that with advances in high-throughput virus whole genome sequencing and the reduction in cost^22^, analysis of this kind will become a powerful tool for understanding the viral genetic variants driving treatment response and whether stratification of patients is required based on viral genetics to optimise treatment outcomes.

Traditionally, RAS have been identified in vivo by comparing baseline sequences to post treatment failure sequences in the protein targeted by the drug^23^. RAS are also identified in-vitro by performing viral selection studies. The majority of RAS have been tested using in vitro assays to determine their impact on viral replication in cell culture^16^. This method has identified sofosbuvir RAS such as NS5B 282T which is treatment emergent but appears transiently due to high fitness cost to the virus^24^. This methodology has two main drawbacks. Firstly, in vitro replicon systems are restricted to a limited number of HCV subtypes and therefore it can be difficult to predict DAA treatment outcome for strains that contain natural polymorphisms which may predispose to reduction in SVR rate^9,10^. Secondly, characterisation of RAS with in vitro approaches has solely focussed on the target protein for a particular class of DAA (e.g. the NS3 gene for DAA compounds that target its protease activity). This ignores the complex interactions between the HCV proteins that are essential to the virus life cycle and the possibility of compensatory mutations in non-target proteins that could also contribute to resistance and treatment failure.

With the benefit of full-length HCV genomic data enabling a comprehensive analysis of all HCV proteins, we have found strong evidence that common polymorphisms in HCV NS2 and NS3 proteins modify treatment outcome in patients receiving SOF+RBV+/-INF. The treatment arm including interferon-alpha had the highest SVR rate and therefore the mechanism of action of the TOPs is unlikely to be associated with interferon-alpha. The mechanism of action of ribavirin is not understood and direct effects on HCV RNA levels are not observed in patients on ribavirin monotherapy^25^, in addition the TOP sites have not been reported in previous studies into ribavirin resistance^26^. Therefore, the most likely mechanism of action of these TOPs is that they reduce the impact of sofosbuvir and therefore affect treatment outcome.

Sofosbuvir targets the NS5B viral RNA polymerase. Although NS2 and NS3 proteins are not directly targeted by sofosbuvir, HCV replication and production is critically dependent on complex interactions between the viral proteins. HCV NS2 is a membrane-bound autoprotease that is required for cleavage at the NS2/NS3 boundary^27–30^. Autoprotease activity is enhanced by the N-terminal region of NS3^27^. Although NS2 is dispensable for HCV RNA replication, the release of NS3 from the polyprotein by NS2 is essential for viral RNA synthesis to proceed. Aside from its autoprotease activity, NS2 plays an essential and central role in assembly of infectious virus, which requires complex interactions with both structural and non-structural proteins^31^. The N-terminal one-third of NS3 also possesses protease activity while the remaining two-thirds encode a RNA helicase^32^. The position of the identified TOPs lie within the protease domains of both proteins^33,34^. In NS2, amino acid position 119 lies in a loop region between two anti-parallel α-helices H1 and H2 while residue 132 is located in α-helix H2^33^. It has been proposed that the loop containing residue 119 may lie close to or interact with a cellular membrane^33^. Position 67 in the NS3 protease domain sits in a loop region between two beta sheets^34^. The functional significance of the variants at these positions in NS2 and NS3 on the virus life cycle is not obvious. Nonetheless, data from in vitro studies have demonstrated that tissue culture adapted variants in HCV proteins that include NS2, NS3 and NS5B can modulate virus production as well as interactions between the viral proteins. Thus, there is the potential for variants to arise which could modify the complex protein-protein interactions needed for both HCV RNA replication and virion assembly, and thereby promote DAA resistance in proteins that are not obvious targets for the drug^35^.

By stratifying the patients, we observed that the reduction in SVR rate based on the combination of these TOPs was broadly additive with patients having both NS2 132V and NS3 67V having an SVR rate of 50% vs. 69% and 86% for patients carrying only NS2 132V or NS3 67V TOP respectively. Patients with viruses that did not carry any TOP had a SVR rate of 94%. We also observed a step-wise reduction in SVR rate as the total number of TOPs present in baseline sequences increased. In studying the impact of these TOPs on reduction in viral load from baseline to week one of therapy, all four TOPs on average resulted in smaller reductions in viral load and this effect was nominally significant for the three sites in NS2 and NS3 proteins. As viral load is a surrogate for viral replication and virion production, its reduction during therapy is an independent measure that can highlight impact on treatment.

We studied the impact of these novel TOPs in an independent cohort of patients treated with sofosbuvir in combination with other DAAs regimes such as NS5A inhibitors (daclatasvir or ledipasvir). The direction and estimated effect sizes were consistent for three of the four sites. However, the associations were not significant as the confidence intervals were larger. It is important to note that this is not an equivalent replication cohort due to its small size (144 patients) and the fact that majority of patients were treated with other DAA in combination with sofosbuvir, which will reduce the power for detecting associations. Additionally, therapy including sofosbuvir as the only DAA is no longer recommended and a cohort similar to BOSON is unlikely to be available in the future.

In conclusion, our data show that common HCV gt3a polymorphisms in NS2 and NS3 proteins are associated with sofosbuvir treatment outcome. Assessing these polymorphisms may be useful in directing therapy length and combinations for difficult to treat patients such as those with advanced cirrhosis or those who have failed DAA therapy previously. Patients carrying both NS2 132V and NS3 67V TOPs had the lowest SVR rate (50%) and the smallest reduction in viral load during the first week of therapy. Patients with these polymorphisms may require special attention if treated using sofosbuvir containing therapies. Sofosbuvir is only used in combination with NS5A inhibitors for which RAS are common in baseline HCV sequences. In such cases polymorphisms that reduces the impact of sofosbuvir will be important in deciding the length and the combination of drugs especially in difficult to treat groups.

## Methods

### BOSON Clinical Trial and Samples

Samples were obtained from patients enrolled in the BOSON clinical trial ^18^ at baseline, during treatment and 12 weeks post treatment. The patients were randomised into three treatment arms of 12 Weeks SOF + RBV + IFN, 16 weeks SOF + RBV and 24 weeks SOF + RBV. All patients were DAA treatment-naïve. Each patient’s HCV viral load was measured at baseline, week 1, week 2, week 4 and week 8. All patients provided written informed consent before undertaking any study-related procedures. The BOSON study protocol was approved by each institution’s review board or ethics committee before study initiation. The study was conducted in accordance with the International Conference on Harmonization Good Clinical Practice Guidelines ^36^ and the Declaration of Helsinki.

### Viral Sequencing

RNA was extracted from 500 μl of plasma using the NucliSENS^®^ easyMAG system (bioMérieux) into 30 μl of water, of which 5 μl was processed with the NEBNext® Ultra™ Directional RNA Library Prep Kit for Illumina® (New England Biolabs) with previously published modifications to the manufacturers protocol ^37^. A 500ng aliquot of the pooled library was enriched using the xGen® Lockdown® protocol from IDT (Rapid Protocol for DNA Probe Hybridization and Target Capture Using an Illumina TruSeq® Library (v1.0), Integrated DNA Technologies) with a comprehensive panel of HCV-specific 120 nucleotide DNA oligonucleotide probes (IDT), designed using a previously published algorithm^38^. The enriched library was sequenced using Illumina MiSeq v2 chemistry to produce paired 150b reads. Reads were demultiplexed low quality reads were trimmed with QUASR and adapter sequences removed using Cutadapt. Host derived sequences were removed using Bowtie. HCV reads were selected by mapping against the 162 ICTV (International Committee on the Taxonomy of Viruses) reference sequences for mapping against the closest reference and de novo assembly (Vicuna), read mapping (MOSAIK), genome annotation (VFAT) and interpretation of variants (genewise2, Vphaser, Vprofiler).

### Statistical Methods

All statistical analysis was performed using the statistical package R, all the analysis code and data is available on request from the STOP-HCV consortium (www.stop-hcv.ox.ac.uk).

#### Non-viral factors associated with outcome

To ensure we accounted for possible confounders when investigating viral effects on outcome, a multivariate analysis was performed using logistic regression to identify non-viral factors, which are associated with SVR. The patient’s cirrhosis status, IFNL4 genotype, prior treatment status, gender, log of baseline viral load and age at time of treatment were investigated as possible confounders. Akaike Information Criterion was used to choose the set of covariates to include in the model. The resulting model (**Supplementary Figure *1***) included, patient’s cirrhosis status, IFNL4 genotype, prior treatment status, gender and log of baseline viral load as covariates in subsequent regression-based analysis unless otherwise stated.

#### Previously reported sofosbuvir RAS effect on outcome

The effect of previously reported sofosbuvir RAS present before treatment was investigated using logistic regression. The response variable was treatment outcome (SVR), the explanatory variable was the presence of RAS and the previously discussed non-viral factors associated with outcome were included as covariates along with the first three viral sequence and host genotyping principle components.

#### Phylogenetics and association of a phenotype with tree structure

RAxML^39^ was used to build a maximum likelihood phylogenetic tree using the HCV full genome sequences assuming the general time reversible model of nucleotide substitution under the gamma model of rate heterogeneity. The treeBreaker^21^ software was then used to investigate if SVR or any novel TOP are associated with specific clades on the tree. If any clades were associated with any of the novel TOP, then the logistic regression was repeated adding an indicator variable for the samples under the clade as a covariate, to confirm that the virus population structure was not confounding the association.

#### Viral genome wide association study

To investigate the effect of individual viral amino acid polymorphisms on SVR a GWAS was performed using a logistic regression model. In our analysis, SVR status was used as the response variable and the presence and absence of each amino acid as the explanatory variable, the non-viral factors previously indicated were included as covariates. To ensure that host-virus population co-structuring had minimal impact on our analysis, we limited our cohort to patients infected with HCV gt3a. To control for both host and virus population structures we included the first three viral PCs calculated from viral sequences and the first three host PCs calculated from the host genome-wide genotyping data as covariates in our model.

#### Replication Cohort

The results were replicated in an independent cohort. Using whole HCV genome sequences from 144 patients enrolled in the HCV Research UK cohort^40^ who were treated with SOF containing regimens. Our hypothesis was that the identified TOPs were associated with outcome therefore one-sided tests were used to replicate the results from the BOSON cohort. Treatment outcome was the response variable and presence/absence of the TOP the explanatory variable. All patients were enrolled in HCVRUK with written, informed consent, all the patients were cirrhotic, and the host genotyping data was not available, so no covariate were included.

#### Covariation and interaction of TOPs analysis

Logistic regression was used to test for association between the number of TOPs in a patient’s viral sequence and SVR. The response variable was SVR and the explanatory variable was number of TOPs in the sequence, the non-viral factors previously indicated were included as covariates. We used logistic regression to investigate possible covariation of TOPs and the first 3 viral PCs and *IFNL4* genotype (CC/non-CC) were added as cofounders as *IFNL4* has been identified as having an effect on the viral genome. Interaction was tested using logistic regression using interaction terms and the confounders identified in **Supplementary Figure 1**.

#### Initial response to therapy

To test if any TOP were associated with a smaller reduction in VL at the start of therapy. The Log10 reduction in viral load from BL to week1 of therapy was calculated. Wilcoxon tests were used to test if patients with TOPs had a lower reduction in viral load than those who did not. Linear regression with the confounders identified in **Supplementary Figure *1*** included was used to test for an association between the number of TOPs in a sequences and reduction in VL.

#### Enrichment of an amino acid in the post-treatment HCV sequences

To test for enrichment of an amino acid in the post-treatment samples (that did not achieve SVR), we used a one-sided binomial test. For each amino acid, the frequency of the amino acid in the baseline sequences, the total number of post-treatment sequences and the observed number of the amino acid in the post-treatment sequences was used to test for its enrichment in the post-treatment sequences.

## Abbreviations

HCV: Hepatitis C Virus
DAA(s): Direct Acting Antivirals
SVR: Sustained Virologic Response
RAS(s): Resistance Associated Substitutions
TOP(s): Treatment Outcome associated Polymorphisms
NS: Non-structural
gt: Genotype
GWAS: Genome Wide Association Study
FDR: False Discovery Rate
IFN: Interferon
IFNL4: Interferon Lambda 4
SNP: Single nucleotide polymorphism

## Supplementary Tables

**Supplementary Table 1:** Top 5 most associated amino acids with SVR in Viral GWAS in order of p-values. We only report the most associated amino acid at each site. At 15% FDR, the first three sites are significantly associated with SVR.

| Position on polyprotein relative to H77 | Protein | Position in Protein and Amino Acid | Available sequence data at position | SVR Rate (%) | Log odds ratio | Standard Error | P value | Q value |
| --- | --- | --- | --- | --- | --- | --- | --- | --- |
| 941 | NS2 | 132V | 506 | 66% | -1.27 | 0.29 | 1.05E-05 | 9.65E-03 |
| 1093 | NS3 | 67V | 506 | 77% | -1.17 | 0.33 | 4.71E-04 | 1.31E-01 |
| 928 | NS2 | 119A | 505 | 59% | -1.88 | 0.54 | 5.18E-04 | 1.31E-01 |
| 1944 | NS4B | 233V | 506 | 58% | 1.54 | 0.50 | 2.01E-03 | 2.95E-01 |
| 2570 | NS5B | 150V | 501 | 75% | -0.76 | 0.27 | 4.92E-03 | 6.21E-01 |

**Supplementary Table 2:** Baseline frequency and SVR rate of amino acids at TOP sites. Only amino acids detected in more than 20 patients are included.

| Protein | Position in Protein | Amino Acid | Frequency at Baseline % | SVR Rate % (n) |
| --- | --- | --- | --- | --- |
| NS2 | 132 | I | 80.0% | 86% (348/405) |
|  |  | V | 19.8% | 66% (66/100) |
| NS3 | 67 | A | 53.6% | 86% (232/271) |
|  |  | V | 43.9% | 77% (171/222) |
| NS2 | 119 | A | 4.4% | 59% (13/22) |
|  |  | M | 4.2% | 81% (17/21) |
|  |  | V | 90.5% | 83% (381/457) |
| NS5B | 150 | A | 38.1% | 88% (168/191) |
|  |  | T | 12.6% | 83% (52/63) |
|  |  | I | 5% | 81% (22/27) |
|  |  | V | 40.7% | 75% (153/204) |

## Supplementary Figures

**Supplementary Figure 1:**
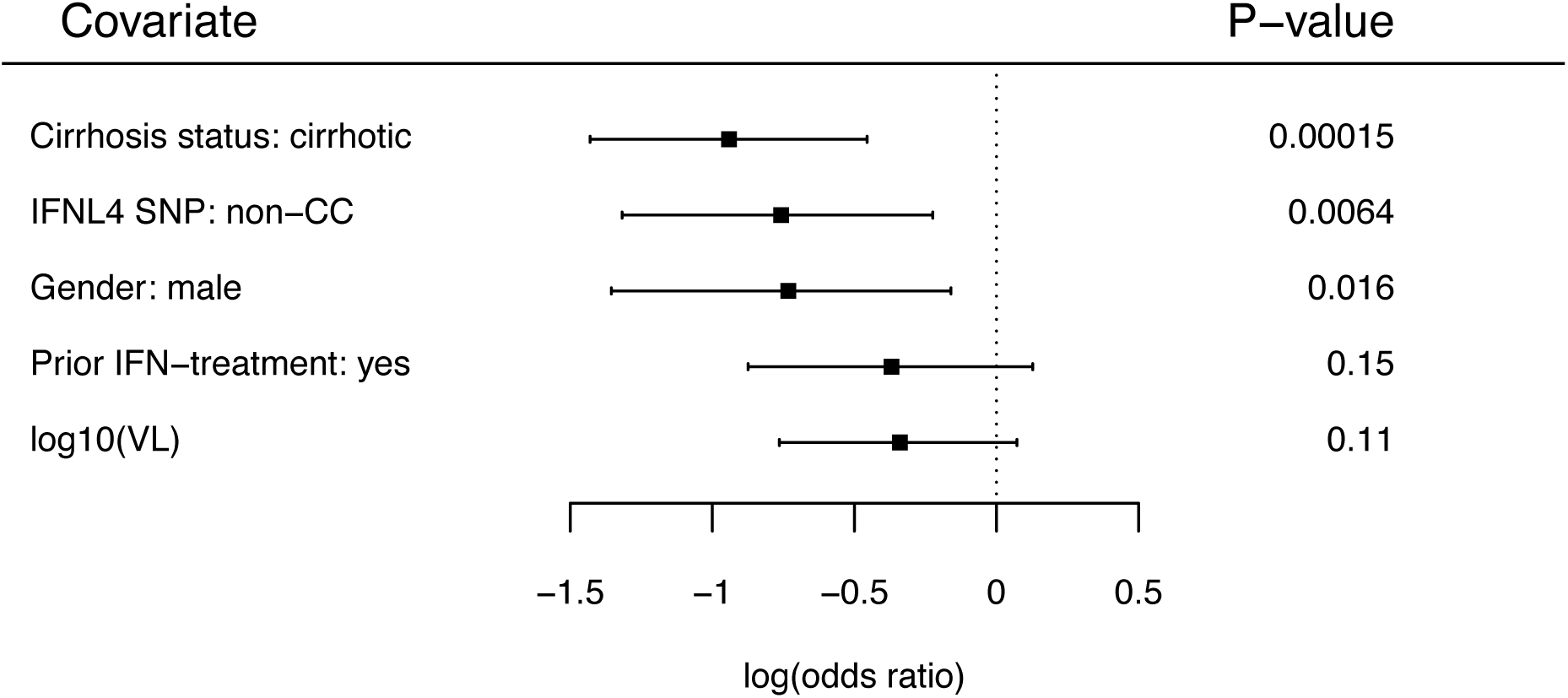
Non-viral factors impact on SVR using a multivariate logistic regression. The squares show the estimated effect size (log(OR)) for each covariate and the lines show its 95% confidence interval, the p value is shown on the right. For the IFNL4 SNP rs12979860 we tested non-CC genotypes (TT and CT) against the CC genotype.

**Supplementary Figure 2:**
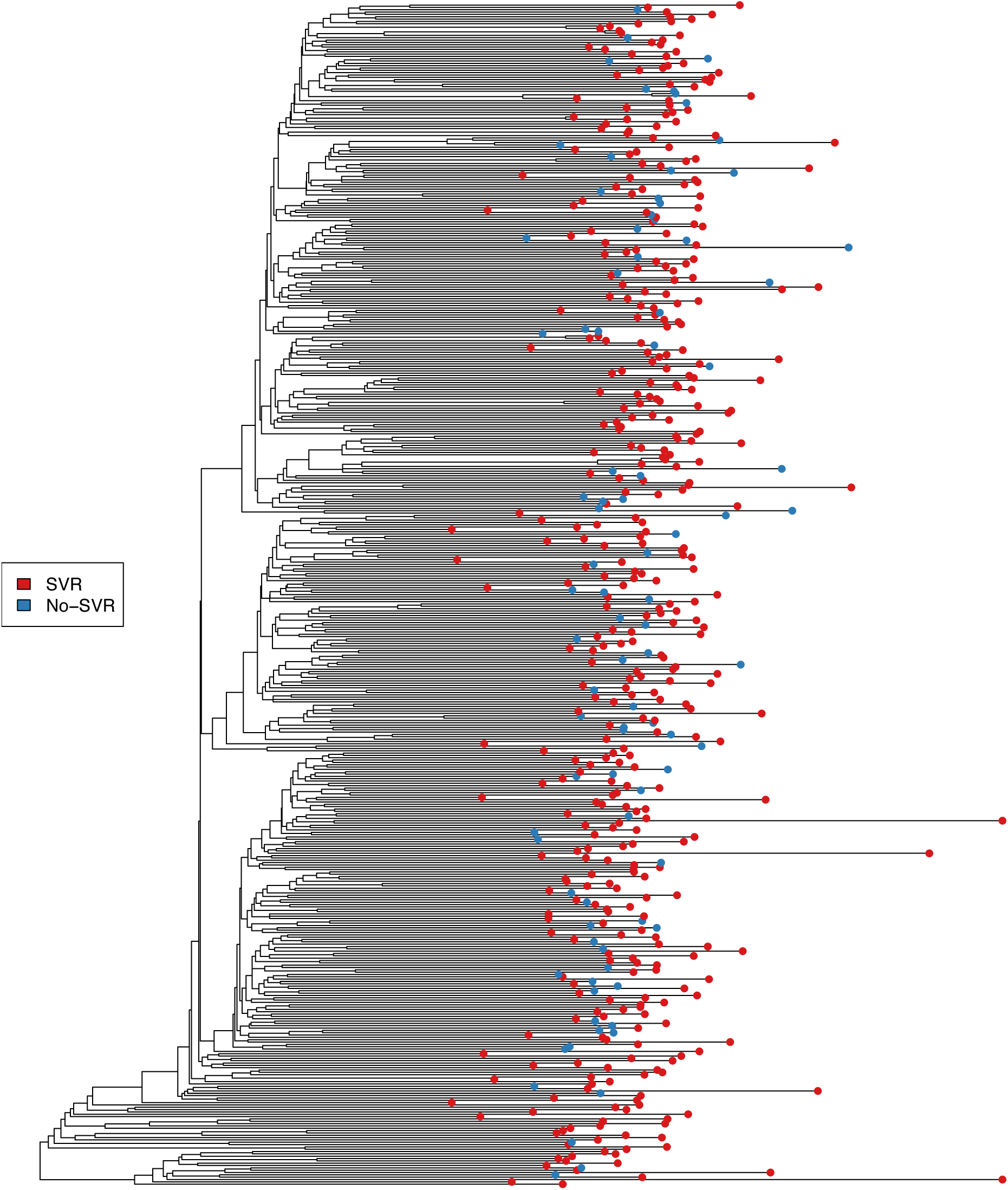
Association between virus population structure and SVR. A maximum likelihood phylogenetic tree for the HCV gt3a sequences was constructed using RAXML. The colour of the tips indicate if the patient achieved SVR (red) or not (blue). The treeBreaker software did not find evidence for any clade to have a different SVR rate from the rest of the tree (Bayes factor = 0.965).

**Supplementary Figure 3:**
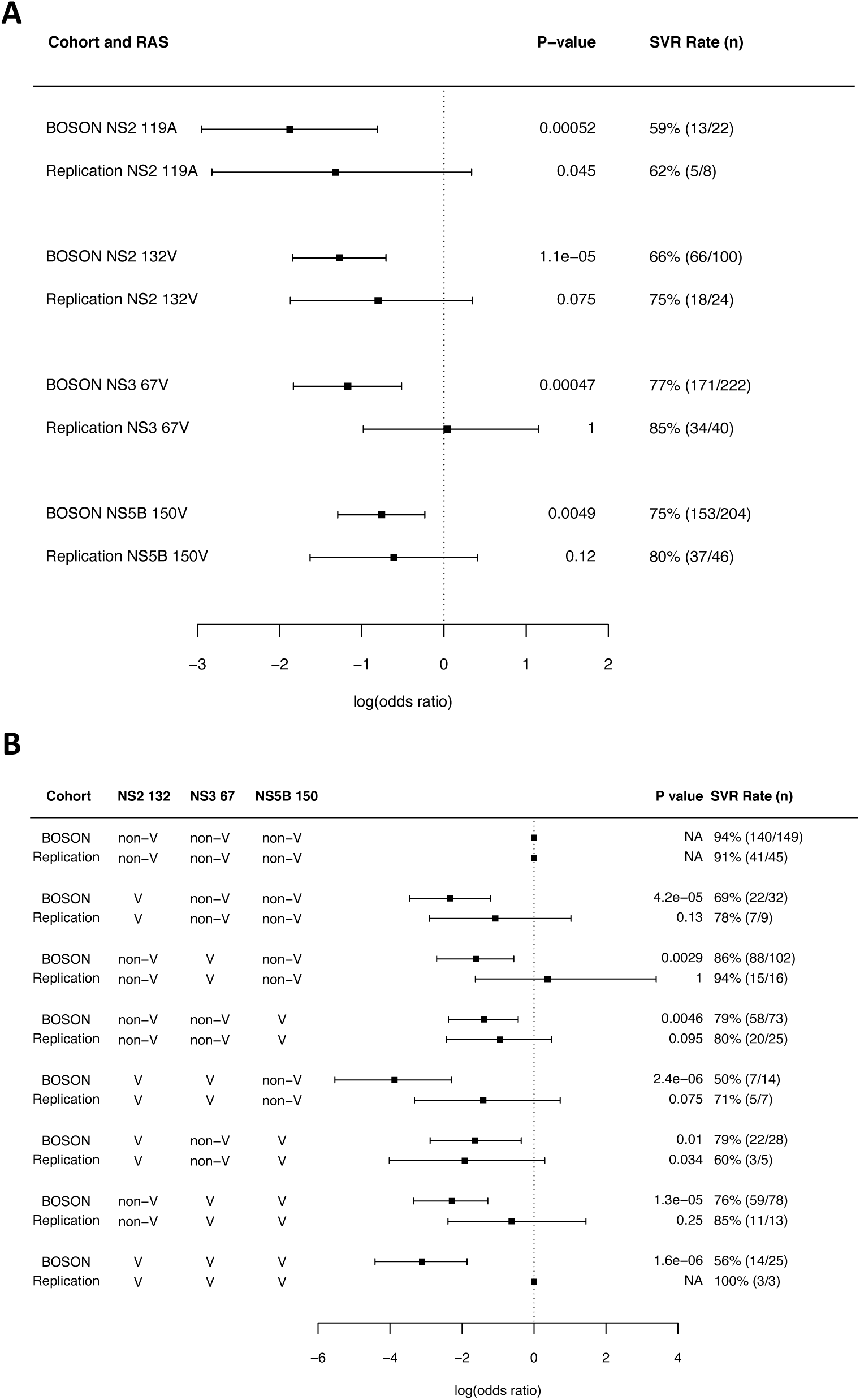
**A) Association between the presence of TOP and outcome in BOSON and replication cohorts.** The cohort and TOP are shown on the right. **B) Covariation between the 3 putative TOP combinations and their association with outcome in both the BOSON and replication cohort.** Combinations tested are listed in the table on the left. The squares show the estimated effect size (log(OR)) for each variable and the lines show its 95% confidence interval. The P values of the associations and SVR rates are shown on the right, P values for the replication cohort are one-sided.

**Supplementary Figure 4:**
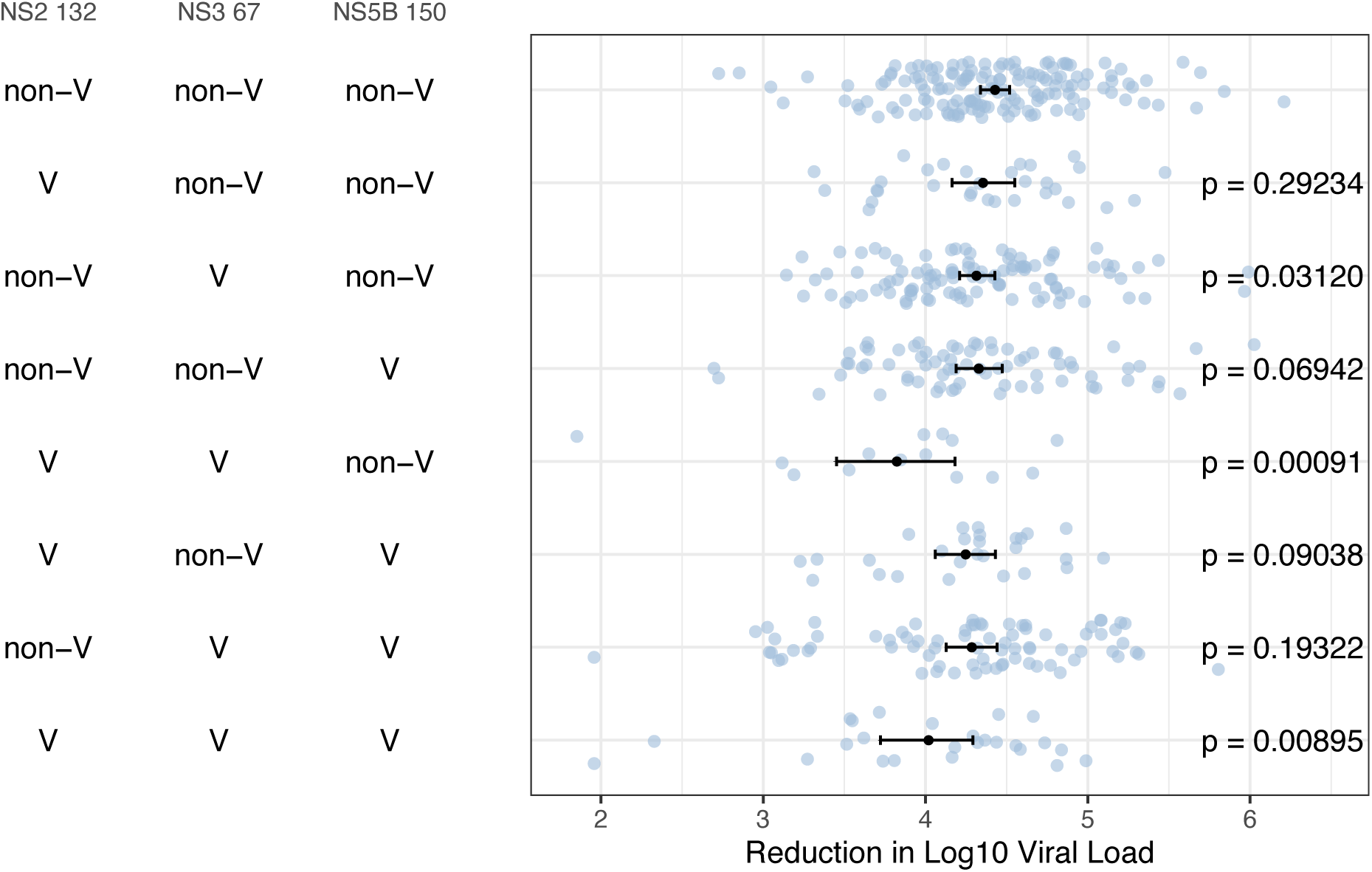
Reduction in log10 viral load between baseline and week 1 of therapy for viruses carrying different combinations of TOP. The mean change in viral load is shown as a black dot for each TOP and the lines indicate its 95% confidence intervals. P values for difference in mean (relative to the no TOP group) calculated using a one sided Mann-Whitney test.

## Notes

**Conflicts of Interest:** GRF: Grants Consulting and Speaker/Advisory Board: AbbVie, Alcura, Bristol-Myers Squibb, Gilead, Janssen, GlaxoSmithKline, Merck, Roche, Springbank, Idenix, Tekmira, Novartis. WLI: Grants, Consulting and Advisory/ Speaker Board: Roche, Janssen Cilag, Gilead Sciences, Novartis, GlaxoSmithKline, Pfizer, Abbvie and Bristol-Myers Squibb. All other authors have no conflicts of interest to declare.

**Financial support statement:** This work was funded by a grant from the Medical Research Council (MR/K01532X/1– STOP-HCV Consortium). The work was supported by Core funding to the Wellcome Trust Centre for Human Genetics provided by the Wellcome Trust (090532/Z/09/Z). EB was/is funded by the Medical Research Council UK, the Oxford NIHR Biomedical Research Centre and is an NIHR Senior Investigator. The views expressed in this article are those of the author and not necessarily those of the NHS, the NIHR, or the Department of Health.

### Competing Interest Statement

GRF: Grants Consulting and Speaker/Advisory Board: AbbVie, Alcura, Bristol-Myers Squibb, Gilead, Janssen, GlaxoSmithKline, Merck, Roche, Springbank, Idenix, Tekmira, Novartis. WLI: Grants, Consulting and Advisory/ Speaker Board: Roche, Janssen Cilag, Gilead Sciences, Novartis, GlaxoSmithKline, Pfizer, Abbvie and Bristol-Myers Squibb. All other authors have no conflicts of interest to declare.

## References

1. World Health Organization. Global hepatitis report, 2017. (World Health Organization, 2017). doi:ISBN 978-92-4-156545-5

2. Feld, J. J. et al. Sofosbuvir and Velpatasvir for HCV Genotype 1, 2, 4, 5, and 6 Infection. N. Engl. J. Med. 373, 2599–607 (2015).

3. Wyles, D. et al. Glecaprevir/Pibrentasvir for HCV Genotype 3 Patients with Cirrhosis and/or Prior Treatment Experience: A Partially Randomized Phase III Clinical Trial. Hepatology (2017). doi:10.1002/hep.29541

4. Niu, C. et al. PSI-7851, a Pronucleotide of -D-2’-Deoxy-2’-Fluoro-2’-C-Methyluridine Monophosphate, Is a Potent and Pan-Genotype Inhibitor of Hepatitis C Virus Replication. Antimicrob. Agents Chemother. 54, 3187–3196 (2010).

5. Zeuzem, S. et al. Sofosbuvir and Ribavirin in HCV Genotypes 2 and 3. N. Engl. J. Med. 370, 1993–2001 (2014).

6. Pawlotsky, J. M. et al. EASL Recommendations on Treatment of Hepatitis C 2018. J. Hepatol. 69, 461–511 (2018).

7. American Association for The Study of Liver Diseases. Recommendations for Testing, Managing, and Treating Hepatitis C.

8. Svarovskaia, E. S. et al. Infrequent development of resistance in genotype 1-6 hepatitis c virus-infected subjects treated with sofosbuvir in phase 2 and 3 clinical trials. Clin. Infect. Dis. 59, (2014).

9. Fourati, S. et al. Frequent antiviral treatment failures in patients infected with hepatitis C virus genotype 4, subtype 4r. Hepatology (2018). doi:10.1002/hep.30225

10. da Silva Filipe, A. et al. Response to DAA therapy in the NHS England Early Access Programme for rare HCV subtypes from low and middle income countries. J. Hepatol. 67, 1348–1350 (2017).

11. Smith, D. et al. Resistance analysis of genotype 3 HCV indicates subtypes inherently resistant to NS5A inhibitors. Hepatology 00, 1–12 (2018).

12. Wei, L. et al. Sofosbuvir–velpatasvir for treatment of chronic hepatitis C virus infection in Asia: a single-arm, open-label, phase 3 trial. Lancet Gastroenterol. Hepatol. (2019). doi:10.1016/S2468-1253(18)30343-1

13. Welzel, T. M. et al. Global epidemiology of HCV subtypes and resistance-associated substitutions evaluated by sequencing-based subtype analyses. J. Hepatol. 67, 224–236 (2017).

14. Naranbhai, V., Hill, A. V. S., Karim, S. S. A. & Fletcher, H. L159F and V321A Sofosbuvir Resistance Associated HCV NS5B Substitutions Accepted. 1–25 (2013). doi:10.1080/08959420.2014.983349

15. Gane, E. J. et al. The emergence of NS5B resistance associated substitution S282T after sofosbuvir-based treatment. Hepatol. Commun. (2017). doi:10.1002/hep4.1060

16. Dietz, J. et al. Patterns of Resistance-Associated Substitutions in Patients With Chronic HCV Infection Following Treatment With Direct-Acting Antivirals. Gastroenterology 154, 976-988.e4 (2018).

17. Wing, P. A. C. et al. Amino Acid Substitutions in Genotype 3a Hepatitis C Virus Polymerase Protein Affect Responses to Sofosbuvir. Gastroenterology (2019). doi:10.1053/j.gastro.2019.05.007

18. Foster, G. R. et al. Efficacy of Sofosbuvir Plus Ribavirin With or Without Peginterferon-Alfa in Patients With Hepatitis C Virus Genotype 3 Infection and Treatment-Experienced Patients With Cirrhosis and Hepatitis C Virus Genotype 2 Infection. Gastroenterology 149, 1462–1470 (2015).

19. Ansari, M. A. et al. Genome-to-genome analysis highlights the effect of the human innate and adaptive immune systems on the hepatitis C virus. Nat Genet advance on, (2017).

20. Tong, X. et al. In vivo emergence of a novel mutant L159F/L320F in the NS5B polymerase confers low-level resistance to the HCV polymerase inhibitors mericitabine and sofosbuvir. J. Infect. Dis. 209, (2014).

21. Azim Ansari, M. & Didelot, X. Bayesian inference of the evolution of a phenotype distribution on a phylogenetic tree. Genetics 204, 89–98 (2016).

22. Cuypers, L. et al. Next-generation sequencing for the clinical management of hepatitis C virus infections: Does one test fits all purposes? Crit. Rev. Clin. Lab. Sci. 1–15 (2019). doi:10.1080/10408363.2019.1637394

23. Donaldson, E. F., Harrington, P. R., O’Rear, J. J., Naeger, L. K. & O’Rear, J. J. Clinical evidence and bioinformatics characterization of potential hepatitis C virus resistance pathways for Sofosbuvir. Hepatology n/a-n/a (2014). doi:10.1002/hep.27375

24. Wyles, D. et al. Post-treatment resistance analysis of hepatitis C virus from phase II and III clinical trials of ledipasvir/sofosbuvir. J. Hepatol. 66, (2017).

25. EASL Clinical Practice Guidelines: management of hepatitis C virus infection. J. Hepatol. 60, 392–420 (2014).

26. Feigelstock, D. A., Mihalik, K. B. & Feinstone, S. M. Selection of hepatitis C virus resistant to ribavirin. Virol. J. 8, 402 (2011).

27. Schregel, V., Jacobi, S., Penin, F. & Tautz, N. Hepatitis C virus NS2 is a protease stimulated by cofactor domains in NS3. Proc. Natl. Acad. Sci. U. S. A. 106, 5342–5347 (2009).

28. Grakoui, A., McCourt, D. W., Wychowski, C., Feinstone, S. M. & Rice, C. M. A second hepatitis C virus-encoded proteinase. Proc. Natl. Acad. Sci. U. S. A. 90, 10583–10587 (1993).

29. Lange, C. M. et al. Determinants for Membrane Association of the Hepatitis C Virus NS2 Protease Domain. J. Virol. 88, 6519–6523 (2014).

30. Hijikata, M. et al. Two distinct proteinase activities required for the processing of a putative nonstructural precursor protein of hepatitis C virus. J. Virol. 67, 4665–4675 (1993).

31. Popescu, C. I. et al. NS2 protein of hepatitis C virus interacts with structural and non-structural proteins towards virus assembly. PLoS Pathog. 7, (2011).

32. Morikawa, K. et al. Nonstructural protein 3-4A: the Swiss army knife of hepatitis C virus. J. Viral Hepat. 18, 305–315 (2011).

33. Lorenz, I. C., Marcotrigiano, J., Dentzer, T. G. & Rice, C. M. Structure of the catalytic domain of the hepatitis C virus NS2-3 protease. Nature 442, 831–835 (2006).

34. Prongay, A. J. et al. Discovery of the HCV NS3/4A protease inhibitor (1R,5S)-N-[3-amino-1-(cyclobutylmethyl)-2,3-dioxopropyl]-3-[2(S)-[[[(1,1-dimethylethyl)amino] carbonyl]amino]-3,3-dimethyl-1-oxobutyl]-6,6-dimethyl-3-azabicyclo[3.1.0] hexan-2(S)-carboxamide (Sch 503034) II. J. Med. Chem. 50, 2310–2318 (2007).

35. Jiang, J. & Luo, G. Cell Culture-Adaptive Mutations Promote Viral Protein-Protein Interactions and Morphogenesis of Infectious Hepatitis C Virus. J. Virol. 86, 8987–8997 (2012).

36. ICH Harmonised Tripartite Guideline. Guideline for good clinical practice E6(R1). ICH Harmon. Tripart. Guidel. 1996, i–53 (1996).

37. Batty, E. M. et al. A Modified RNA-Seq Approach for Whole Genome Sequencing of RNA Viruses from Faecal and Blood Samples. PLoS One 8, (2013).

38. Bonsall, D. et al. ve-SEQ: Robust, unbiased enrichment for streamlined detection and whole-genome sequencing of HCV and other highly diverse pathogens. F1000Research 4, 1062 (2015).

39. Stamatakis, A. RAxML version 8: A tool for phylogenetic analysis and post-analysis of large phylogenies. Bioinformatics 30, 1312–1313 (2014).

40. McLauchlan, J. et al. Cohort Profile: The Hepatitis C Virus (HCV) Research UK Clinical Database and Biobank. Int. J. Epidemiol. 46, (2017).

